# *dMARCH8,* a *Drosophila* ubiquitin E3 ligase, regulates polymodifications of tubulin in the spermiogenic pathway

**DOI:** 10.1101/112243

**Authors:** Ujwala A. S. Gosavi, Nabanita Chatterjee, Utsav Nyachhyon, Brian R Pearce, Robert J Harvey, Christopher Bazinet

**Affiliations:** Department of Biological Sciences, St. John’s University, Jamaica, New York 11439, USA; Department of Pharmacology, UCL School of Pharmacy, 29-39 Brunswick Square, London WC1N 1AX, UK

## Abstract

Ciliary stability and function are regulated by the covalent addition of polyglutamate and polyglycine chains to axonemal tubulin subunits. The *Drosophila* gene CG13442 encodes a predicted ubiquitin E3 ligase involved in the regulation of tubulin glycylation and glutamylation. Homologous to mammalian MARCH8, CG13442/*dMARCH8* is required for male fertility. Sperm in *dMARCH8* mutant testes appear to undergo a normal individualization process but fail to be transferred to the seminal vesicle. This phenotype is very similar to that of mutants in the *Ntl* gene, shown here to be a glycine transporter using a [^3^H]glycine uptake assay. Mutations in *dMARCH8* are associated with a reduction of both polyglutamylation and polyglycylation of sperm tubulin. Polyglutamylation of tubulin is significantly increased in the *Ntl*^−^ background, and recovers to wild-type levels in the *Ntl*^−^-dMARCH8^−^ double mutant background, indicating that glycine and glutamate compete for some common site(s) on tubulin molecules in this system. By analogy to the regulation of the mammalian glycine transporter GlyT2 through ubiquitin-mediated trafficking between the plasma membrane and endosome, *dMARCH8* may target *Ntl* and glutamate transporters, or other upstream regulators of these proteins.

## Introduction

Amino acids and their chemical derivatives serve as signaling molecules in several biological contexts. Glycine and glutamate function as major inhibitory and excitatory neurotransmitters, respectively, in the central nervous system (Kandel et al., 2013). At the subcellular level, these same two amino acids regulate ciliary function and stability through a mechanism involving their competition for covalent addition of polyglycine and polyglutamate chains to tubulin subunits incorporated into stable microtubule-based structures (Raunser and Gatsogiannis, 2015). The activity of glycine transporters, controlling the levels of glycine both inside and outside of the cell, can therefore result in behavioral effects and ciliary dysfunction (Boison, 2016)

Here we use the *Drosophila* spermiogenesis system to examine the regulation of tubulin polymodifications. *Drosophila* spermiogenesis provides an opportunity for genetic analysis of many different cellular subsystems (Fabian and Brill, 2012). These include the differentiation of mitochondria (Hales and Fuller, 1997;Politi et al., 2014), polarization of the spermiogenic cyst (Wei et al., 2008), the radical restructuring of spermatid nuclei (Kost et al., 2015), and the complex process of sperm individualization, in which individual sperm are finally resolved from the syncytium in which they have developed (Arama et al., 2007;Arama et al., 2003;Fabrizio et al., 1998;Tokuyasu et al., 1977) This complexity is reflected in the large number of genes that are mutable to a male-sterile phenotype, and indicates that genes contributing to any of the many different cellular subsystems---including the polymodifications of tubulin---may be accessible through the analysis of male-sterile mutations (Wakimoto et al, 2007).

Because there is very little transcription in spermiogenic cysts (Barreau et al., 2008) posttranscriptional and posttranslational regulatory mechanisms are especially important in this process (Karr, 2007). Ubiquitination is one of these processes, affecting many different aspects of spermiogenesis (Richburg et al., 2014). Although the ubiquitin system is known primarily for marking proteins for degradation (Ciechanover, 2005;Zhi et al., 2013), recent studies indicate that ubiquitination can affect many other aspects of protein function, including intracellular trafficking (de Juan-Sanz et al., 2011), modulation of protein-protein interactions (Yang et al., 2010), and modulation of transcription, DNA repair and transmembrane signaling (Metzger et al., 2012).

The substrate-specificity of a ubiquitination system is generally controlled by E3 ubiquitin ligases. Here we identify an E3 ubiquitin ligase, *dMARCH8,* homologous to mammalian MARCH8. *dMARCH8/CG13442* function is required for the full complement of polyglycylation and polyglutamylation of sperm tubulin in the *Drosophila* testis. We show that the *Drosophila Neurotransmitter transporter-like (Ntl)* gene, previously found to exhibit a similar phenotype to *dMARCH8* mutants (Chatterjee et al., 2011), encodes a glycine transporter using a [^3^H]glycine uptake assay. Loss of *Ntl* function results in a large increase in polyglutamylation levels, indicating that *Ntl* and *dMARCH8* activities contribute to the balance of glutamylation and glycylation of sperm tubulin.

## Materials and Methods

### Fly Husbandry

Flies were raised on standard cornmeal molasses agar at 25°C. Unless otherwise mentioned, all stocks were from the Bloomington Stock Center. Males were tested for fertility by mating in groups of 4–5 with an equal number of virgin females. Generally, *w*^+^ or *y*^+^ males were mated with *yw* females, with the recovery of *w*^+^ or *y*^+^ daughters in the F1 generation confirming fertility. Genetic constructions were carried out using standard *Drosophila* genetics as in Greenspan (Greenspan RJ, 1997). All experiments were carried out in the *yw* genetic background.

### Generation *of dMARCH8* mutants

The *dMARCH8* transcript/CDR is in the 57B region on the 2R arm of the *Drosophila* chromosome. The Mi{MIC}(*Minos* Mediated Integrated Cassette) *dpr^MI06571^* transposon carrying the *y^+^* marker was generated by the *Drosophila* Gene disruption project (GDP)(Venken et al., 2011)The transposon was mobilized by crossing the flies carrying the MiMIC insertion to flies carrying stable heat shock transposase (*y*^1^ *w*\*; *sna^Sco^ / SM6a, P{w^+mC^=hsILMiT}2.4* (Metaxakis et al., 2005;Venken et al., 2011). Three broods from each cross were generated by transferring flies to new bottles on the 3^rd^ and 5^th^ days after the cross was initially set up. Bottles were subjected to heat shock (39°C, 90 minutes) on the third, 5^th^ and seventh day of each brood. Chromosomes that lost the *y*^+^ markers carried by the transposon were recovered and established in balanced stocks using standard *Drosophila* genetics, then screened for new male-sterile mutations produced by imprecise excision of the Mi{MIC} transposon.

### RNA isolation and qRT-PCR

Total RNA was isolated using TRI reagent (Sigma) according to the manufacturer’s recommendations. RNA was extracted from males, females, testes, and heads and concentration was determined by measuring its absorbance.

One-step qRT-PCR was performed using the iTaq Universal SYBR Green One-Step Kit (BIORAD) according to the manufacturer’s recommendations. The BIORAD MyiQ Real-Time PCR Detection System was programmed as follows: 50°C for 10 min, 95°C for 5 min followed by the amplification steps of 95°C for 15 sec, 56°C for 30 seconds, 72°C for 30 sec. 45 cycles of PCR were run for all samples followed by 72°C for 15 min, 57°C for 10 min and held at 4°C overnight. *dMARCH8* qRT primers amplified a 112 bp fragment, while rp49 control primers amplified a 154 bp fragment. Primers for this and all subsequent PCR-based molecular biology are specified in Table 1.

**Table 1:**
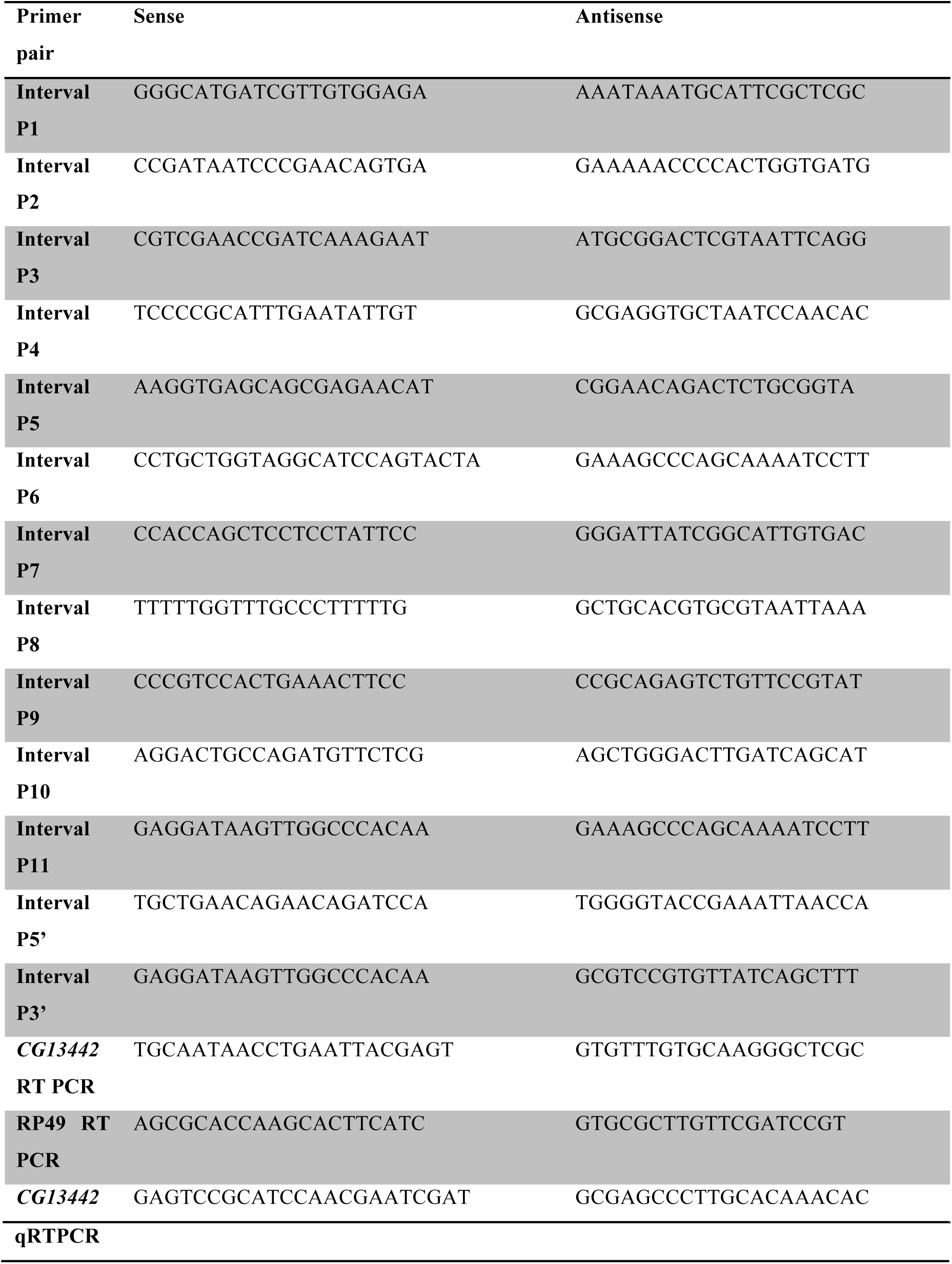
Primers.

### Deletion PCR

Genomic DNA was isolated from males according to the Berkeley *Drosophila* Genome Project protocol (http://www.fruitfly.org/about/methods/inverse.pcr.html). 13 pairs of gene-specific primers were used to amplify the entire coding sequence of *dMARCH8*. The thermocycler (MJ Research PTC-200 Peltier thermal cycler) was programmed as follows: 94°C for 2 min followed by 30 cycles of: 94°C for 1 min, 56.3°C for 1 min, 72°C for 1 min. After a final 5 min at 72°C, samples were held at 4°C until gel analysis.

### Phalloidin assay for Individualization Complex

Testes from 0–1 d old males were crudely dissected in *Drosophila* Ringers and transferred immediately to a tube of Ringers on ice. Testes were then fixed for 15 min in 4% paraformaldehyde in buffer B (16.7mM KH_2_PO_4_/K_2_HPO_4_ pH 6.8, 75mM KCl, 25mM NaCl, 3.3mM MgCl_2_). Following fixation, testes were rinsed three times in PTx (PBS+0.1% Triton X-100), washed for 30 min in PTx and blocked for at least 1 h in blocking solution (0.01% NaAzide and 3% BSA in PTx). Staining with rhodamine-conjugated phalloidin was for 30 min in blocking solution (3 μg/ml). Testes were then rinsed and washed with PTx. Testes were then finely dissected from remaining carcasses in 50% glycerol and then mounted in 90% glycerol. Slides were stored overnight at 4°C before imaging. Fluorescence images were captured by confocal microscopy (Leica TCS-2, Exton, PA)(Fabrizio et al., 2012)

### Phase squashes

Testes were dissected from 0–1-day-old young male *Drosophila* mutants (unless mentioned otherwise) in 1 × PBS buffer and gently squashed with a coverslip before taking phase images with Leica DM 4500B. For phase microscopy of seminal vesicles, wild type and mutant freshly eclosed *Drosophila* males were withheld from females for three to four days before dissecting their testes.

### Protein Electrophoresis and Immunoblot

Samples were prepared from male fly testes with seminal vesicle for each genotype, from males that were withheld from females for 4–6 days in order to maximize sperm yield. Six fly testes (12 testes) worth of protein were loaded in each lane. Testes were dissected in 1 × PBS, ground in 2× Laemmeli buffer (Laemmli UK., 1970) vortexed and boiled for 5 min. Samples were then spun at 15,800 g for 5 min, and supernatants were separated in a 12% SDS-Polyacrylamide gel and transferred to a PVDF membrane (Amersham, GE Healthcare,United Kingdom) using a Trans-Blot Semi-Dry transfer apparatus (BioRad, U.S.A). Membranes were incubated with primary antibodies directed against poly-glycylated tubulin (Poly-G) (1:10,00) (MABS276, cloneAXO49, Millipore), polyglutamylated tubulin (Poly-E) (1:1000) (mAbGT355, Adipogen Life sciences), ubiquitin {MAB1510 clone Ubi-1 (aka 042691GS), Millipore}, TSSK2 (ab172434, Abcam) and α-tubulin (1:500), (DM1A, Sigma). Protein bands were visualized with HRP-labelled anti rabbit or anti-mouse (1:5,000) secondary antibodies followed by detection with ECL immunoblot detection kit (Pierce, U.S.A). Loading control was α-tubulin.

All statistical analyses were made using two-tailed Students t-test in Microsoft Excel. The average values of relative intensity (Target antibody intensity/anti-tubulin) were plotted for each genotype. Intensities were calculated using Image J software (Rasband, W.S., ImageJ, U. S. National Institutes of Health, Bethesda, Maryland, USA, http://rsb.info.nih.gov/ij/, 1997–2009).

### [3H]glycine uptake assays

Human GlyT1, GlyT2 and *Drosophila* Ntl cDNAs were cloned into pRK5myc or pTMR vectors as previously described (Carta et al., 2012). HEK293 cells were grown in minimal essential medium (Earle’s salts) supplemented with 10% (v/v) FCS, 2 mm L-glutamine, and 20 units/ml penicillin/streptomycin in 5% CO_2_, 95% air. The cells were plated on poly-D-lysine-coated 24-wells plates (Nunc), grown to 50% confluence, and transfected with 1 μg of total pRK5myc-hGlyT1, pRK5myc-hGlyT2, pRK5myc-Ntl or pTMR-Ntl plasmid DNAs using Lipofectamine LTX reagent (Invitrogen). After 24 h, the cells were washed twice with prewarmed buffer (118 mM NaCl, 1 mM NH_2_PO_4_, 26 mM NaHCO_3_, 1.5 mM MgSO_4_, 5 mM KCl, 1.3 mM CaCl_2_, 20 mM glucose) pre-equilibrated with 5% CO_2_, 95% air. After 2 min, the cells were incubated for 5 min in 0.1 μCi/ml [^3^H]glycine (60 Ci/mmol; PerkinElmer Life Sciences) at a final concentration of 300 μM. The cells were rinsed twice with ice-cold buffer pre-equilibrated with 5% CO_2_, 95% air and then digested in 0.1 M NaOH for 2 h. The samples were used for scintillation counting and for determination of protein concentration using the Bradford reagent (Bio-Rad). [^3^H]glycine uptake was calculated as nmol/min/mg of protein and expressed as percentages of that in control cells transfected with the empty expression vector. All statistical comparisons used an unpaired Student’s *t*-test.

### Generation of *Ntl-dMARCH8* double mutant

*Ntl-dMARCH8* double mutants were generated by crossing *Ntl/CyO* males to *dMARCH8^7A^/CyO* females. Straight-winged female progeny obtained were crossed to *yw*, *Gla/Sm6a* males. Isomales from this cross were crossed again to establish balanced stocks. Males from these established stocks were checked for fertility by crossing to *yw* female virgins. Males from sterile stocks obtained were screened for complementation with *Ntl*^−^ and *dMARCH8*^−^ mutants.

## Results

### Amino acid sequence alignment of *dMARCH8 (CG13442)* with human MARCH8-E3 ubiquitin Ligases belonging to the RING protein family

A ClustalW alignment of the predicted dMARCH8 protein sequence with human MARCH8 is shown in Figure 1A. RING domain considered critical for E3 ubiquitin ligase function (Samji et al., 2014) are conserved in the predicted dMARCH8 protein. The TMpred algorithm (Hofmann and Stoffel, 1993) predicts two transmembrane domains for dMARCH8, characteristic of the MARCH family (Samji et al., 2014), whose RING domains are most closely related to the Ringv and PHD domains (Deshaies and Joazeiro, 2009).

**Figure 1:**
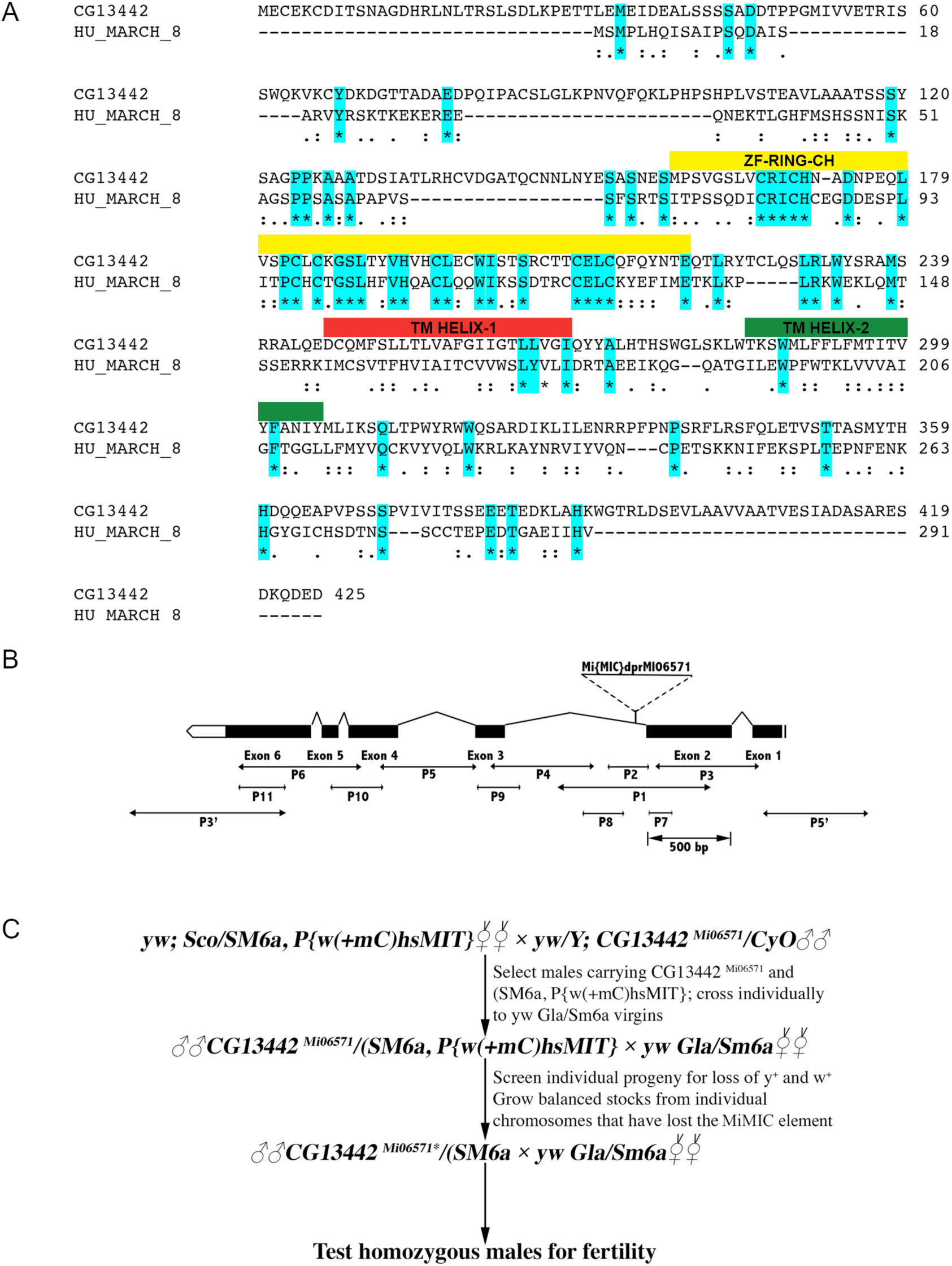
Amino acid sequence alignment of *Drosophila melanogaster CG13442 (dMARCH8)* with human MARCH 8 using ClustalW alignment and generation of mutants in the *dMARCH8* (CG13442) gene. A) Strictly conserved residues are highlighted in blue. Conserved RING-CH domain is highlighted in yellow and transmembrane domains 1 and 2 are highlighted in red and green respectively. B) *dMARCH8* (CG13442) transcript/CDR at 57B1 on the right arm of chromosome 2. MiMIC transposon element is inserted between Exon2 and Exon3, closer to Exon2. C) Scheme for generation of *dMARCH8* (CG13442) deletion mutations by imprecise excision of the MiMIC transposon

### Generation of *dMARCH8* mutants

The *dMARCH8* gene lies within an intron of the *dpr* gene, where it is expressed from the opposite strand. *dMARCH8* mutants were generated by mobilizing the Mi{MIC}*dpr^MI06571^* near the 5’ end of the *dMARCH8* gene, crossing it into a genetic background expressing the minos transposase (Venken et al., 2011) under heat shock control (Metaxakis et al., 2005). By standard fly genetics, chromosomes which had lost the *yellow*^+^ (*y*^+^) markers associated with the Mi{MIC} transposon were recovered and established in balanced stocks. Transposon excisions are often imprecise and deletions of varying size, usually extending from the insertion site in either direction, are often recovered at significant frequencies (Zhang and Spradling, 1993). A schematic of the *dMARCH8* locus showing the starting insertion, the mating scheme used to mobilize the transposon and identify new male-sterile mutations, and the location of primer pairs used to assay for deletions in the resulting male-sterile stocks is shown in Figure 1B and 1C. Of 64 chromosomes that were observed to have lost the *y*^+^ marker, 5 were found to carry new male sterile mutations defining a single complementation group.

### Deletion PCR analysis of *CG13442/dMARCH8* mutants

The mutant alleles were screened with nested gene-specific primers spanning the site of the transposon insertion (Materials and Methods and Fig. 1B). Genomic DNA from homozygous males carrying each of the 5 male-sterile alleles was used to probe for changes in the chromosome structure by PCR. For all of the male-sterile mutants, PCR amplicons including portions of the gene and flanking regions on either side were missing or produced products with altered size as compared to wild-type DNA (Table 2). These results indicated that the deletion carried by the *dMARCH8^7A^* mutant removes the sequences encoding the RING domain and both transmembrane domains (Table 2). Furthermore, qRT-PCR analysis of this mutant revealed a significant reduction (80 +/− 5% in *dMARCH8* expression (Fig. 2A). The *dMARCH8^7A^* allele was used for the remaining phenotypic analyses.

**Table 2:**
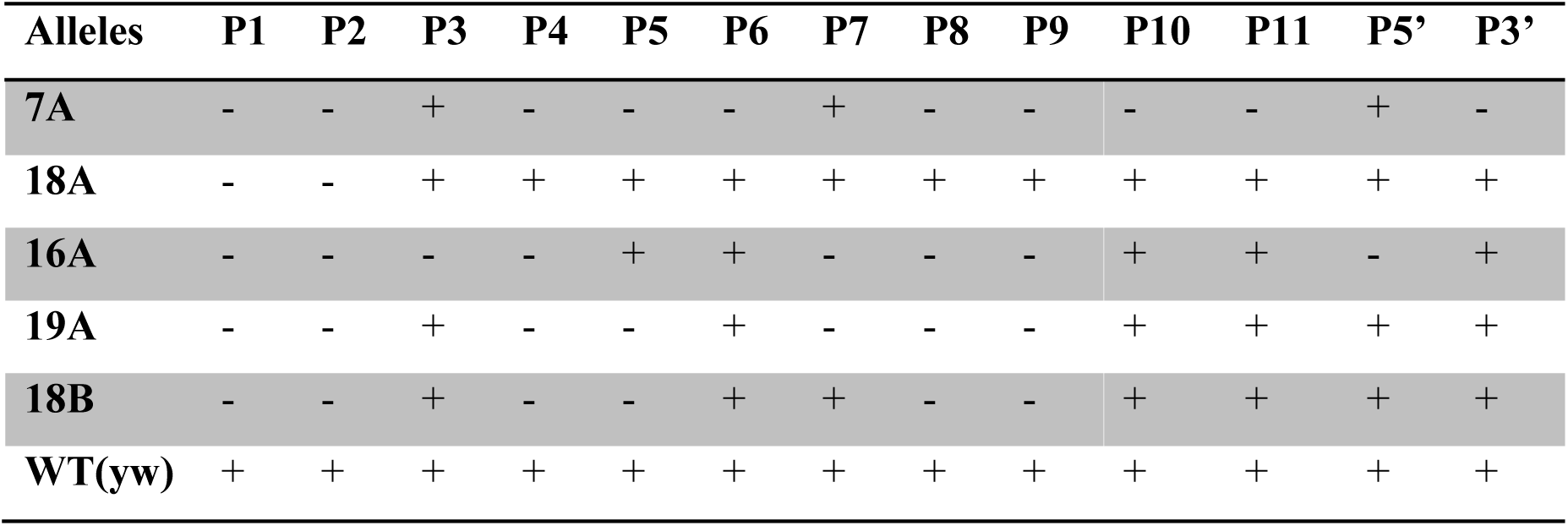
Deletion Analysis of CG13442 mutants. +/− Refer to presence or absence of the band respectively

**Figure 2:**
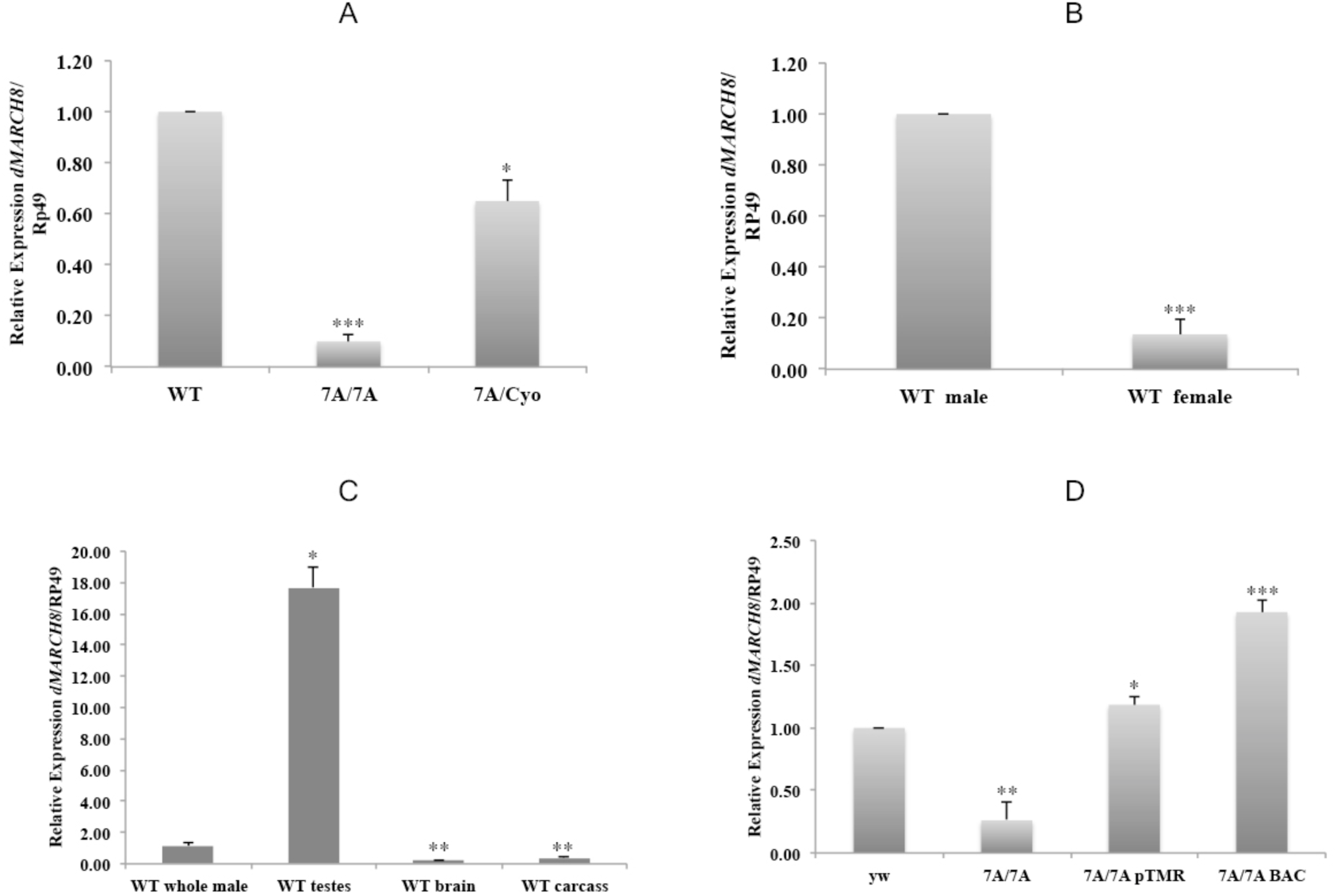
Analysis of *dMARCH8* mRNA Expression. A) qRT-PCR analysis shows decrease in *dMARCH8* expression levels in *dMARCH8^7A^/dMARCH8^7A^* (7A/7A) mutants as compared to wild-type(*yw*) and heterozygous *dMARCH8^7A^/CyO* flies. Bar, standard error of the mean. N=3 (* p<0.05, ** p<0.01, **p<0.001 by two-tailed t-test). B) qRT-PCR analysis compares the relative expression of *dMARCH8* in wild-type (*yw*) male and female (whole adult flies). Bar, standard error of the mean. N=3 (* p<0.05, ** p<0.01, **p<0.001 by two-tailed t-test). C) qRT-PCR products from wild-type (*yw*) whole male, (*yw*) male testes, (*yw*) male brain, and (*yw*) male carcass. Bar, standard error of the mean. N=3 (* p<0.05, ** p<0.01, **p<0.001 by two-tailed t-test). D) *dMARCH8* qRT-PCR products from *dMARCH8^7A^/dMARCH8^7A^* (7A/7A) mutant males carrying a pTMR-*dMARCH8* cDNA construct (*yw*; *dMARCH8^7A^/dMARCH8^7A^*; pTMR-*dMARCH8*) (7A/7A;pTMR) and a genomic BAC construct (*yw*; *dMARCH8^7A^/dMARCH8^7A^*; BAC*-dMARCH8*) (7A/7A;BAC) compared to mutant *dMARCH8^7A^/dMARCH8^7A^* and wild-type controls (*yw*). Bar, standard error of the mean. N=3 (* p<0.05, ** p<0.01, **p<0.001 by two-tailed t-test).

### Testis Specificity of *dMARCH8* expression

qRT-PCR analysis confirmed that *dMARCH8* expression was male-specific and limited to the testes (Fig. 2B and C). Expression in females was 90 +/− 5 % lower than in males, and in the adult heads and carcass of males, it was 98 +/− 1% lower than in the testis.

### Rescue by Germline Transformation

We used a P-element based construct for germline transformation with CG13442/ *dMARCH8*, to confirm that the male sterile phenotype in our mutants is caused solely due to disruption of *dMARCH8*. For this we used a pTMR-*dMARCH8* construct, which contains a full-length cDNA for CG13442 (DGRC clone AT03090) cloned downstream of the β2T-tubulin transcriptional control sequences in the *Drosophila* transformation vector pTMR (Clark et al., 2006;Huh et al., 2004). This construct provides strong germ cell-specific transcription in developing sperm (Kaltschmidt et al., 1991;Kemphues et al., 1982). A BAC genomic construct CH322-140N02 (BACPAC resources, bacpac.chori.org) extending from 9.8 kb upstream of the *dMARCH8* transcript to 8.5 kb beyond the 3’ end also rescued the *dMARCH8* mutant phenotype. Two independent insertions of this construct were tested and they both rescued the mutant. All the males homozygous for the *dMARCH8^7A^* deletion and carrying pTMR-*dMARCH8 or BAC-dMARCH8* constructs were fertile. Results from the rescued line are presented in Fig. 2D. The male-sterile mutations obtained after mobilization of the Mi{MIC} transposon therefore result from disruption of *dMARCH8*.

### Spermiogenic-defective phenotype of *dMARCH8* mutants

*dMARCH8^7A^* / *dMARCH8^7A^* males produced elongated spermiogenic cysts (Fig. 3), but we did not detect mature sperm in the seminal vesicle (SV) (Fig. 3B, arrows). In contrast, wild-type control seminal vesicles were filled with mature sperm (Fig. 3A, arrows). No motile sperm are seen in *dMARCH8^7A^* / *dMARCH8^7A^* squash preparations, unlike in the wild-type controls, where dense masses of mature motile sperm were evident (Fig. 3C, arrow). Instead, an extensive mass of sperm bundles accumulated at the base of *dMARCH8^7A^* mutant testes (Fig. 3B, asterisks). Presumably because of the great physical complexity of the sperm individualization process (Bazinet and Rollins, 2003;Noguchi et al., 2008;Tokuyasu et al., 1972), a preponderance of male-sterile mutations produce elongated cysts that fail to mature into individual sperm (Wakimoto et al, 2007). In most of these cases, the individualization complex either fails to form, does not progress, or breaks down during transfer along the length of the cyst (Fabrizio et al., 1998).

**Figure 3:**
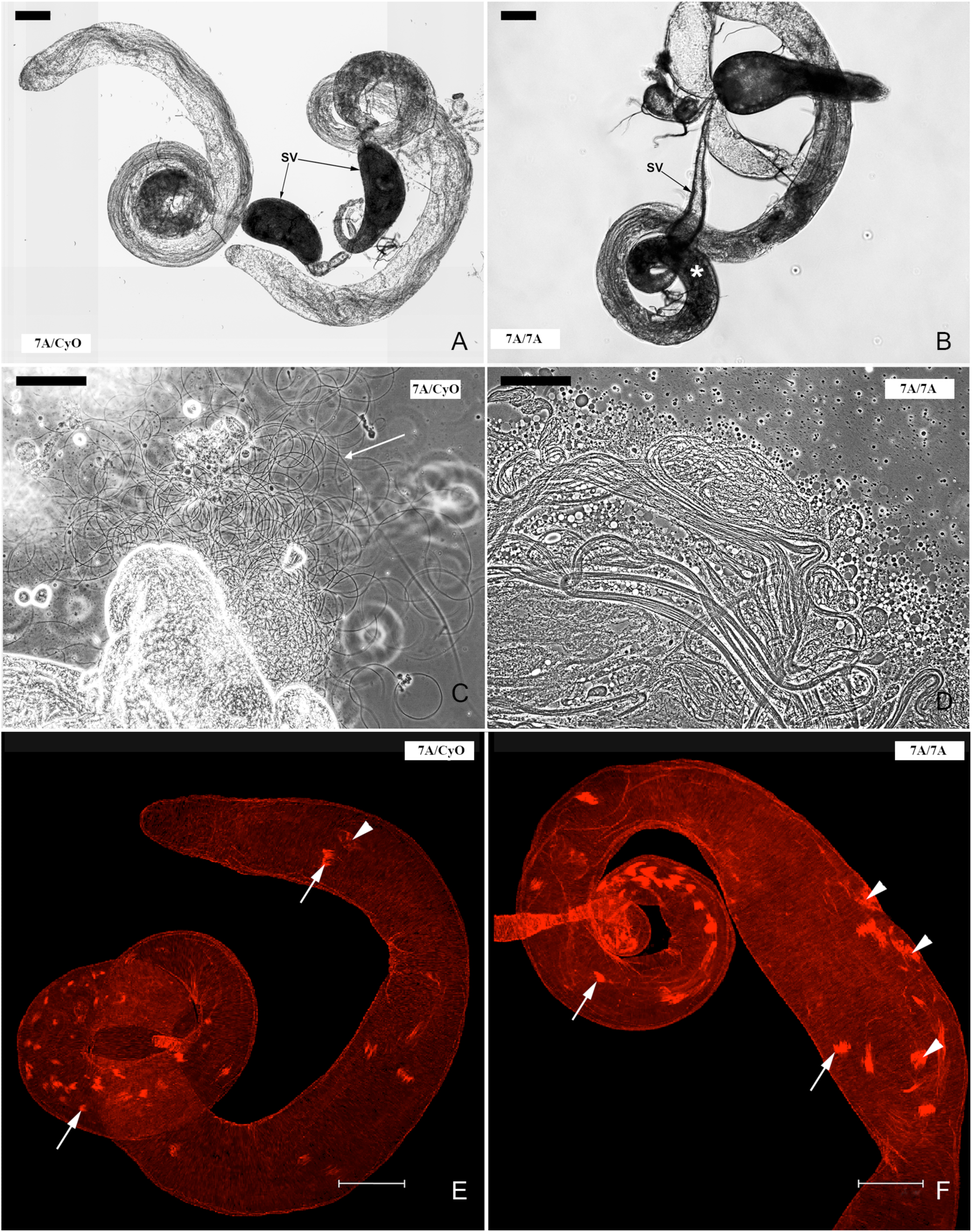
Immotility of *dMARCH8* mutant sperm, failing to transfer into seminal vesicles and individualization in *dMARCH8* mutants. Panels A, B, C and D: Phase contrast images of testes from *dMARCH8^+^* (A, C) and *dMARCH8*^−^ (B, D) males. The major phenotypic feature of the mutants is the accumulation of coiled cysts at the base of the testis (asterisks), and the empty/shrunken state of the seminal vesicle (SV) (arrows). In panel C, the letter M denotes dense masses of mature motile sperm, which is not seen in the mutants. Left hand panels: heterozygous *(dMARCH8^7A^/Cyo)*; right hand panels: *(dMARCH8^7A^/dMARCH8^7A^)* mutants. Bars, 100 μm. E: Heterozygous *dMARCH8^7A^/Cyo* testis stained with rhodamine-conjugated phalloidin to visualize the actin cone-based individualization complexes. Arrows mark the actin cones of the complex. Arrowheads mark the waste-bags. F: *dMARCH8^7A^/ dMARCH8^7A^* mutant testis preparations stained with rhodamine-conjugated phalloidin to visualize the actin cone-based individualization complexes. Formation and movement of actin cones/individualization complex along the mutant cysts appears normal. Bars, 100 um.

When *dMARCH8^7A^* mutant *Drosophila* testes were stained with rhodamine-conjugated phalloidin, we observed normal development and movement of the actin cones of the individualization complex (Fig. 3F). Waste bag deposition in the distal end of the testis also appeared normal, further indicating that the individualization complex was successfully navigating the entire length of the cysts (Fig 3F). These observations are similar to the phenotype seen in *Neurotransmitter transporter-like* (*Ntl*) mutants, a putative glycine transporter homologous to human glycine transporters GlyT1 and GlyT2 (Chatterjee et al., 2011).

### Demonstration of Glycine transport activity from Ntl expression in mammalian cells

To test whether the *Drosophila* Ntl protein was capable of glycine uptake, we tested the capacity of recombinantly-expressed Ntl to mediate the uptake of [^3^H]glycine, using human GlyT1 and GlyT2 cDNAs (Carta et al., 2012) as controls. As expected, both pRK5-GlyT1 and pRK5-GlyT2 constructs resulted in significant increases in [^3^H]glycine uptake compared to vector only controls (pRK5-GlyT1: 345 ± 42; pRK5-GlyT2 299 ± 38). Both Drosophila Ntl expression constructs also resulted in statistically significant [^3^H]glycine uptake (pRK5-NTL: 245 ± 24; pTMR-NTL: 344 ± 59) confirming that Ntl is a fully-functional glycine transporter (Fig. 4).

**Figure 4:**
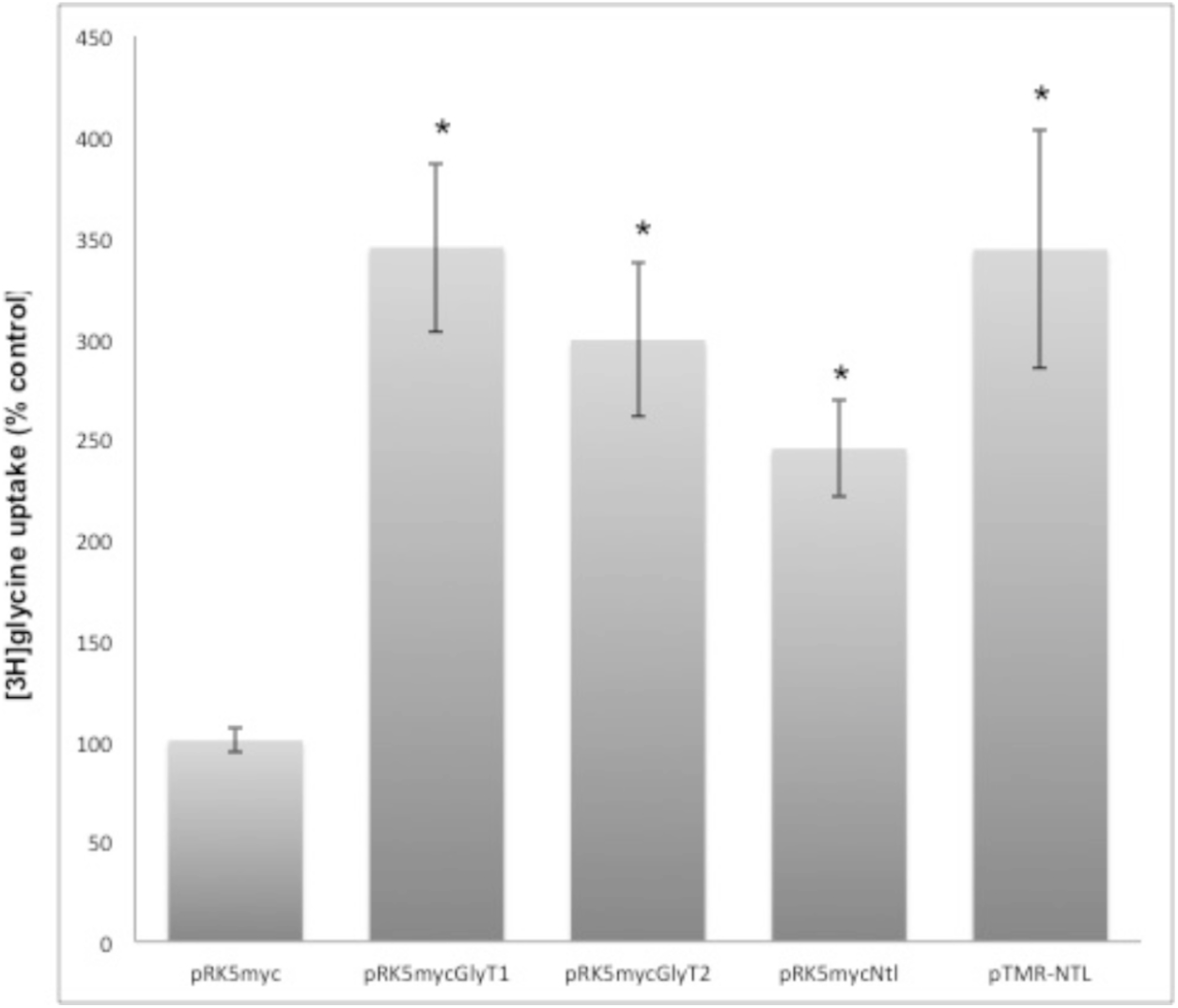
Drosophila Ntl is a functional glycine transporter. Glycine uptake in HEK293 cells transiently expressing hGlyT1, hGlyT2 and *Drosophila* Ntl expressed from two different expression vectors after 5 min of incubation with [^3^H]glycine at a final concentration of 300 μM. Because low levels of glycine uptake are found in HEK293 cells (Carta et al., 2012), [^3^H]glycine uptake was calculated as nmol/min/mg of protein and then expressed as a percentage of the empty expression vector (pRK5-myc) transfected control. The data are the means ± S.E. (*n* = 7-8). Statistical comparisons were made using an unpaired Students *t*-test. The *asterisk* indicates significantly different from empty vector control (*p* < 0.01).

### Reduction of ubiquitination of testis proteins in *dMARCH8* mutants

Since dMARCH8 is a member of the E3 ubiquitin ligase family, we assayed *dMARCH8* mutant males for total ubiquitination of testis proteins using an anti-ubiquitin antibody (Kane et al., 2014). The results from three independent experiments, using α-tubulin as a loading control, showed an apparent reduction in ubiquitination signal in mutant samples relative to wild-type controls (Fig. 5A). Two distinct ubiquitinated bands of ~50 and 77 kDa present in the testes of *dMARCH8^7A^*/+ flies are missing in the testes of homozygous *dMARCH8^7A^* flies. A slight reduction in several other bands (~65 and 100 kDa) was also consistently observed in the *dMARCH8^7A^* mutant.

**Figure 5:**
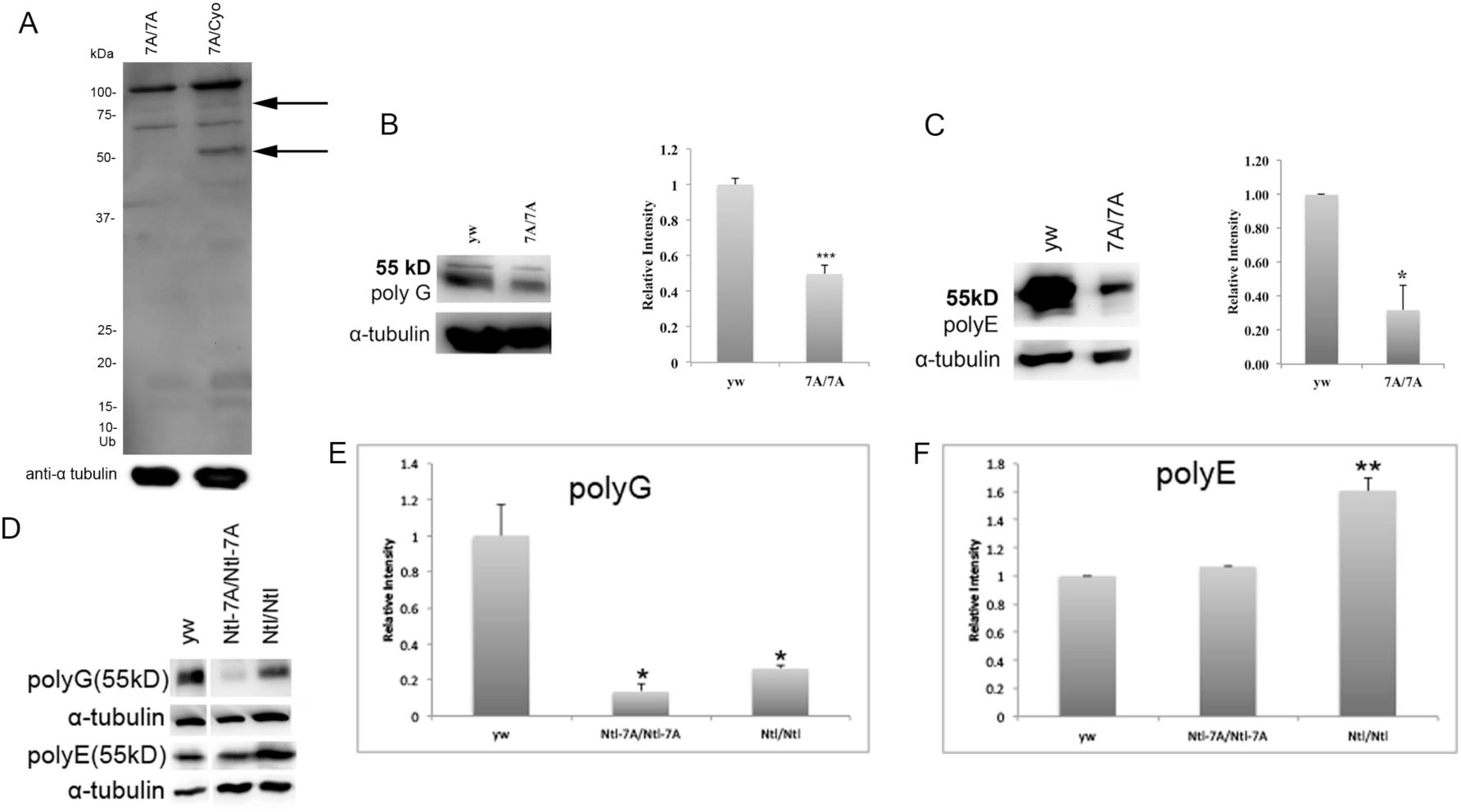
Western Blot Analysis of *dMARCH8^7A^/dMARCH8^7A^, Ntl/Ntl* and *Ntl-dMARCH8^7A^* mutant testes. A) Total ubiquitination levels of testis proteins are reduced in dMARCH8 mutant testes. Western blot of dMARCH87A/dMARCH87A (dMARCH8-) males compared to dMARCH87A/CyO (dMARCH8+) heterozygous males probed with anti-ubiquitin antibody. B) Polyglycylation of tubulin is partially decreased in mutant testes. Western blot of dMARCH87A/dMARCH87A (dMARCH8-) males compared to yw (dMARCH8+/+) probed with anti Poly-G antiserum. Quantitation of three independent replicates of the Western analysis. Bar, standard error of the mean. N=3 (* p<0.05, ** p<0.01, **p<0.001 by two-tailed t-test). C) Polyglutamylation of tubulin is partially decreased in mutant testes. Western blot of dMARCH87A/dMARCH87A (dMARCH8-) males compared to yw (dMARCH8+/+) controls probed with anti Poly-E antiserum. Quantitation of three independent replicates of the Western analysis. Bar, standard error of the mean. N=3 (* p<0.05, ** p<0.01, **p<0.001 by two-tailed t-test). (D, E, F) Polyglycylation (polyG) and polyglutamylation (polyE) levels of tubulin in Ntl/Ntl and Ntl-dMARCH87A mutant testes. Western blot showing tubulin polyglycylation and polyglutamylation levels in Ntl/Ntl (lane 3) and Ntl-dMARCH87A/ Ntl-dMARCH87A (lane 2) mutant testes compared to yw (WT) (lane 1) probed with anti Poly-G and Poly-E antiserum. Quantitation of three independent replicates of the polyG and polyE Western analysis respectively. Bar, standard error of the mean. N=3 (* p<0.05, ** p<0.01, **p<0.001 by two-tailed t-test).

### Reduction of tubulin polyglycylation and polyglutamylation in *dMARCH8* mutants

The tubulin in *Drosophila* sperm axonemes undergoes a variety of posttranslational modifications (Janke, 2014;Kierszenbaum, 2002;Wloga and Gaertig, 2010)Polyglycylation of tubulins is required for the stability of ciliary and flagellar axonemes and other long-lived tubulin-based structures (Bre et al., 1996;Bressac et al., 1995;Rogowski et al., 2009a). Polyglutamylation is known to regulate beating behavior in motile cilia via the regulation of flagellar dynein motors (Ikegami et al., 2010;Janke et al., 2005;Kubo et al., 2010;Pathak et al., 2007;Suryavanshi et al., 2010). In addition, polyglycylation and polyglutamylation have been shown to affect male fertility (Chatterjee et al., 2011;Lee et al., 2013).

The spermiogenic defect of *dMARCH8* mutants at the microscopic level is similar to that seen in *Ntl*, which encodes a glycine transporter in whose absence polyglycylation of testes tubulin is significantly reduced and sperm fail to be transferred to the seminal vesicle. We therefore analyzed the levels of polyglycylated and polyglutamylated tubulin in testis protein samples using antibodies directed against poly-G and poly-E. Quantitation by scanning the results from six independent experiments, using α-tubulin as a loading control showed an average of 50% reduction in poly-G signal in the *dMARCH8^7A^* mutant samples relative to wild-type controls (Fig 5B). Similarly, for the poly-E signal, quantitation from three independent experiments exhibited an average of 70% reduction in *dMARCH8^7A^* mutants compared to wild-type controls (Fig. 5C).

### Polyglycylation and polyglutamylation in a *Ntl-dMARCH8* double mutant

Since the *Ntl* mutant exhibits a similar phenotype to the *dMARCH8* mutant, and it also shows a predicted interaction (STITCH database) with dMARCH8, we analyzed the *Ntl-dMARCH8* double mutant for polyglycylation and polyglutamylation levels of tubulin in three independent experiments. We observed that in both the *Ntl* mutant and the *Ntl-dMARCH8* double mutant, poly-G levels decrease while the poly-E levels increase (Fig. 5D, E and F). Perhaps most strikingly, loss of *Ntl* in the *dMARCH8*^-^ genetic background restores levels of polyglutamylation to wild-type levels (compare Fig. 5C and 5F). In this system, polyglutamylation appears to be very sensitive to glycine levels, consistent with observations by others that glutamate and glycine compete with each other for common site(s) on tubulin (Bulinski, 2009;Rogowski et al., 2009b;Wloga et al., 2009).

## Discussion

Functional spermatogenesis is particularly dependent on ubiquitin-regulated protein function and stability (Mukhopadhyay and Riezman, 2007;Richburg et al., 2014). This is evident from the spermatogenesis-specific functions of a number of E3 ligases: Bruce and Cullin3 in *Drosophila* have been studied for their role in sperm individualization (Arama et al., 2007;Arama et al., 2003;Kaplan et al., 2010;Wang et al., 2006). RNF8 and E3^histone^, found in mice and rats respectively, are reported to be involved in the histone degradation that occurs during histone to protamine transition in spermatid nuclei in rats (Liu et al., 2005;Lu et al., 2010). The E3 ligases Cul4A (Kopanja et al., 2011;Yin et al., 2011), Itch (Dwyer and Richburg, 2012) and Siah1a (Dickins et al., 2002) play roles during germ cell meiosis and Cullin3^testis^ (Arama et al., 2007;Kaplan et al., 2010) and MEX (Nishito et al., 2006) are required during germ cell apoptosis. MARCH7 (<u>M</u>embrane-<u>A</u>ssociated <u>R</u>ing-<u>CH</u>) was found to be involved in spermiogenesis by regulating the structural and functional integrity of the head and tail of developing spermatids (Zhao et al., 2013). MARCH10 was found to be essential for spermatid maturation (Iyengar PV, Hirota T, Hirose S, Nakamura N., 2011). MARCH11 plays a role in ubiquitin-mediated protein sorting in TGN-MVB (Trans Golgi Network-multivesicular bodies) transport in developing spermatids (Morokuma et al., 2007;Yogo et al., 2012).

Indeed, of all organs examined in rats, ubiquitination has been reported to be highest in testis (Rajapurohitam et al., 2002). Combined with the observed genetic sensitivity of the spermiogenic process, these considerations indicate that details of cellular ubiquitin function may be accessible through the characterization of male-sterile mutations in E3 ligase genes. Studies of mammalian spermatogenesis have implicated ubiquitination in the regulation of multiple spermiogenic stages, including nuclear condensation, acrosome formation and membrane transport (Nakamura, 2013).

Here we have observed that loss of the E3 ubiquitin ligase *dMARCH8* results in a male-sterile phenotype very similar to that observed in mutants of the transporter *Ntl.* We have also shown *Ntl* mediates the uptake of [^3^H]glycine in a cellular assay with an efficiency comparable to the mammalian glycine transporters GlyT1 and GlyT2. At least one other uncharacterized *Drosophila* SLC6 class transporter encoded by CG5549, gives a somewhat better BLAST score against human GlyT1 (42.34% identity) and GlyT2 (35.01 % identity) than *Ntl* (Ntl vs GlyT1 39.15%; Ntl vs GlyT2 37.6% identity) suggesting that the fly genome may encode an additional glycine transporter. Expression of CG5549 is also male biased {Flybase}, although not as strongly as that of *Ntl*.

Modulation of the activity of the mammalian glycine transporter GlyT in mammalian cells is thought to be achieved via ubiquitin-dependent endocytosis of GlyT to recycling endosomes, from which it can be returned to the plasma membrane when needed (Barrera et al., 2015;de Juan-Sanz et al., 2011;de Juan-Sanz et al., 2013;Fernandez-Sanchez et al., 2009). Ubiquitylation of the GlyT is associated with internalization of the transporter, where it can where it can no longer contribute to cellular glycine uptake. In that case, one might expect constitutively higher levels of glycine in cells that had lost the ability to ubiquitinate the transporter. If the Ntl transporter was similarly regulated through ubiquitylation by *dMARCH8*, one might expect that loss of *dMARCH8* would not result in reduced availability of glycine as a result of the intracellular sequestration of the Ntl transporter. Instead, the opposite is observed, a reduction in polyglycylation and glutamylation. This suggests that *Ntl* and potentially a glutamate transporter are not themselves substrates of *dMARCH8,* but that the reduced glycylation and glutamylation results from the activity of *dMARCH8* on factors upstream of glycine/glutamate transport, or in some parallel pathway. Future studies will seek to clarify whether Ntl is a substrate of dMARCH8 and to identify and characterize any additional components of the system controlling spermiogenic tubulin glycylation and glutamylation.

## References

Arama, E., Bader, M., Rieckhof, G. E. and Steller, H. (2007). A ubiquitin ligase complex regulates caspase activation during sperm differentiation in drosophila. PLoS Biol. 5, e251.

Arama, E., Agapite, J. and Steller, H. (2003). Caspase activity and a specific cytochrome C are required for sperm differentiation in drosophila. Developmental Cell 4, 687–697.

Barreau, C., Benson, E., Gudmannsdottir, E., Newton, F. and White-Cooper, H. (2008). Post-meiotic transcription in drosophila testes. Development 135, 1897–1902.

Barrera, S. P., Castrejon-Tellez, V., Trinidad, M., Robles-Escajeda, E., Vargas-Medrano, J., Varela-Ramirez, A. and Miranda, M. (2015). PKC-dependent GlyT1 ubiquitination occurs independent of phosphorylation: Inespecificity in lysine selection for ubiquitination. PLoS One. 10, e0138897.

Bazinet, C. and Rollins, J. E. (2003). Rickettsia-like mitochondrial motility in drosophila spermiogenesis. Evol. Dev. 5, 379–385.

Boison, D. (2016). The biochemistry and epigenetics of epilepsy: Focus on adenosine and glycine. Front. Mol. Neurosci. 9, 26.

Bre, M. H., Redeker, V., Quibell, M., Darmanaden-Delorme, J., Bressac, C., Cosson, J., Huitorel, P., Schmitter, J. M., Rossler, J., Johnson, T. et al. (1996). Axonemal tubulin polyglycylation probed with two monoclonal antibodies: Widespread evolutionary distribution, appearance during spermatozoan maturation and possible function in motility. J. Cell. Sci. 109 (Pt 4), 727–738.

Bressac, C., Bre, M. H., Darmanaden-Delorme, J., Laurent, M., Levilliers, N. and Fleury, A. (1995). A massive new posttranslational modification occurs on axonemal tubulin at the final step of spermatogenesis in drosophila. Eur. J. Cell Biol. 67, 346–355.

Bulinski, J. C. (2009). Tubulin posttranslational modifications: A pushmi-pullyu at work? Dev. Cell. 16, 773–774.

Carta, E., Chung, S. K., James, V. M., Robinson, A., Gill, J. L., Remy, N., Vanbellinghen, J. F., Drew, C. J., Cagdas, S., Cameron, D. et al. (2012). Mutations in the GlyT2 gene (SLC6A5) are a second major cause of startle disease. J. Biol. Chem. 287, 28975–28985.

Chatterjee, N., Rollins, J., Mahowald, A. P. and Bazinet, C. (2011). Neurotransmitter transporter-like: A male germline-specific SLC6 transporter required for drosophila spermiogenesis. PLoS One. 6, e16275.

Ciechanover, A. (2005). Proteolysis: From the lysosome to ubiquitin and the proteasome. Nat. Rev. Mol. Cell Biol. 6, 79–87.

Clark, I. E., Dodson, M. W., Jiang, C., Cao, J. H., Huh, J. R., Seol, J. H., Yoo, S. J., Hay, B. A. and Guo, M. (2006). Drosophila pink1 is required for mitochondrial function and interacts genetically with parkin. Nature 441, 1162–1166.

de Juan-Sanz, J., Nunez, E., Lopez-Corcuera, B. and Aragon, C. (2013). Constitutive endocytosis and turnover of the neuronal glycine transporter GlyT2 is dependent on ubiquitination of a C-terminal lysine cluster. PLoS One. 8, e58863.

de Juan-Sanz, J., Zafra, F., Lopez-Corcuera, B. and Aragon, C. (2011). Endocytosis of the neuronal glycine transporter GLYT2: Role of membrane rafts and protein kinase C-dependent ubiquitination. Traffic 12, 1850–1867.

Deshaies, R. J. and Joazeiro, C. A. (2009). RING domain E3 ubiquitin ligases. Annu. Rev. Biochem. 78, 399–434.

Dickins, R. A., Frew, I. J., House, C. M., O’Bryan, M. K., Holloway, A. J., Haviv, I., Traficante, N., de Kretser, D. M. and Bowtell, D. D. (2002). The ubiquitin ligase component Siah1a is required for completion of meiosis I in male mice. Mol. Cell. Biol. 22, 2294–2303.

Dwyer, J. L. and Richburg, J. H. (2012). Age-dependent alterations in spermatogenesis in itchy mice. Spermatogenesis 2, 104–116.

Fabian, L. and Brill, J. A. (2012). Drosophila spermiogenesis: Big things come from little packages. Spermatogenesis 2, 197–212.

Fabrizio, J. J., Aqeel, N., Cote, J., Estevez, J., Jongoy, M., Mangal, V., Tema, W., Rivera, A., Wnukowski, J. and Bencosme, Y. (2012). Mulet (mlt) encodes a tubulin-binding cofactor E-like homolog required for spermatid individualization in drosophila melanogaster. Fly (Austin) 6, 261–272.

Fabrizio, J. J., Hime, G., Lemmon, S. K. and Bazinet, C. (1998). Genetic dissection of sperm individualization in drosophila melanogaster. Development 125, 1833–1843.

Fernandez-Sanchez, E., Martinez-Villarreal, J., Gimenez, C. and Zafra, F. (2009). Constitutive and regulated endocytosis of the glycine transporter GLYT1b is controlled by ubiquitination. J. Biol. Chem. 284, 19482–19492.

Greenspan RJ. (1997). The theory and practice of DrosophilaGenetics. Plainview, new york: Cold spring harbor laboratory press. Fly Pushing.

Hales, K. G. and Fuller, M. T. (1997). Developmentally regulated mitochondrial fusion mediated by a conserved, novel, predicted GTPase. Cell 90, 121–129.

Hofmann, K. and Stoffel, W. (1993). TMBASE - A database of membrane spanning protein segments. Biol Chem Hoppe-Seyler 374,.

Huh, J. R., Vernooy, S. Y., Yu, H., Yan, N., Shi, Y., Guo, M. and Hay, B. A. (2004). Multiple apoptotic caspase cascades are required in nonapoptotic roles for drosophila spermatid individualization. PLoS Biol. 2, E15.

Ikegami, K., Sato, S., Nakamura, K., Ostrowski, L. E. and Setou, M. (2010). Tubulin polyglutamylation is essential for airway ciliary function through the regulation of beating asymmetry. Proc. Natl. Acad. Sci. U. S. A. 107, 10490–10495.

Iyengar PV, Hirota T, Hirose S, Nakamura N. (2011). Membrane-associated RING-CH 10 (MARCH10 protein) is a microtubule-associated E3 ubiquitin ligase of the spermatid flagella. ;286(45):39082–39090. doi:10.1074/jbc.M111.256875. The Journal of Biological Chemistry. 286, 39082–39090.

Janke, C. (2014). The tubulin code: Molecular components, readout mechanisms, and functions. J. Cell Biol. 206, 461–472.

Janke, C., Rogowski, K., Wloga, D., Regnard, C., Kajava, A. V., Strub, J. M., Temurak, N., van Dijk, J., Boucher, D., van Dorsselaer, A. et al. (2005). Tubulin polyglutamylase enzymes are members of the TTL domain protein family. Science 308, 1758–1762.

Kaltschmidt, B., Glatzer, K. H., Michiels, F., Leiss, D. and Renkawitz-Pohl, R. (1991). During drosophila spermatogenesis beta 1, beta 2 and beta 3 tubulin isotypes are cell-type specifically expressed but have the potential to coassemble into the axoneme of transgenic flies. Eur. J. Cell Biol. 54, 110–120.

Kandel, E., Schwartz, J., Jessell, T., Siegelbaum, S. and Hudspeth, A. (2013). Principles of neural science. Fifth Edition, Mc Graw Hill, New York,.

Kane, L. A., Lazarou, M., Fogel, A. I., Li, Y., Yamano, K., Sarraf, S. A., Banerjee, S. and Youle, R. J. (2014). PINK1 phosphorylates ubiquitin to activate parkin E3 ubiquitin ligase activity. J. Cell Biol. 205, 143–153.

Kaplan, Y., Gibbs-Bar, L., Kalifa, Y., Feinstein-Rotkopf, Y. and Arama, E. (2010). Gradients of a ubiquitin E3 ligase inhibitor and a caspase inhibitor determine differentiation or death in spermatids. Developmental Cell 19, 160–173.

Kemphues, K. J., Kaufman, T. C., Raff, R. A. and Raff, E. C. (1982). The testis-specific beta-tubulin subunit in drosophila melanogaster has multiple functions in spermatogenesis. Cell 31, 655–670.

Kierszenbaum, A. L. (2002). Sperm axoneme: A tale of tubulin posttranslation diversity. Mol. Reprod. Dev. 62, 1–3.

Kopanja, D., Roy, N., Stoyanova, T., Hess, R. A., Bagchi, S. and Raychaudhuri, P. (2011). Cul4A is essential for spermatogenesis and male fertility. Dev. Biol. 352, 278–287.

Kost, N., Kaiser, S., Ostwal, Y., Riedel, D., Stützer, A., Nikolov, M., Rathke, C., Renkawitz-Pohl, R. and Fischle, W. (2015). Multimerization of drosophila sperm protein Mst77F causes a unique condensed chromatin structure. Nucleic Acids Research 43, 3033–3045.

Kubo, T., Yanagisawa, H. A., Yagi, T., Hirono, M. and Kamiya, R. (2010). Tubulin polyglutamylation regulates axonemal motility by modulating activities of inner-arm dyneins. Curr. Biol. 20, 441–445.

Laemmli UK. (1970). Cleavage of structural proteins during the assembly of the head of bacteriophage T4. Nature 227, 680–685.

Lee, G. S., He, Y., Dougherty, E. J., Jimenez-Movilla, M., Avella, M., Grullon, S., Sharlin, D. S., Guo, C., Blackford, J. A., Jr, Awasthi, S. et al. (2013). Disruption of Ttll5/stamp gene (tubulin tyrosine ligase-like protein 5/SRC-1 and TIF2-associated modulatory protein gene) in male mice causes sperm malformation and infertility. J. Biol. Chem. 288, 15167–15180.

Liu, Z., Oughtred, R. and Wing, S. S. (2005). Characterization of E3(histone), a novel testis ubiquitin protein ligase which ubiquitinates histones. Molecular and Cellular Biology 25, 2819–2831.

Lu, L., Wu, J., Ye, L., Gavrilina, G. B., Saunders, T. L. and Yu, X. (2010). RNF8-dependent histone modifications regulate nucleosome removal during spermatogenesis. Developmental Cell 18, 371–384.

Metaxakis, A., Oehler, S., Klinakis, A. and Savakis, C. (2005). Minos as a genetic and genomic tool in drosophila melanogaster. Genetics 171, 571–581.

Metzger, M. B., Hristova, V. A. and Weissman, A. M. (2012). HECT and RING finger families of E3 ubiquitin ligases at a glance. J. Cell. Sci. 125, 531–537.

Morokuma, Y., Nakamura, N., Kato, A., Notoya, M., Yamamoto, Y., Sakai, Y., Fukuda, H., Yamashina, S., Hirata, Y. and Hirose, S. (2007). MARCH-XI, a novel transmembrane ubiquitin ligase implicated in ubiquitin-dependent protein sorting in developing spermatids. J. Biol. Chem. 282, 24806–24815.

Mukhopadhyay, D. and Riezman, H. (2007). Proteasome-independent functions of ubiquitin in endocytosis and signaling. Science 315, 201–205.

Nakamura, N. (2013). Ubiquitination regulates the morphogenesis and function of sperm organelles. Cells 2, 732–750.

Nishito, Y., Hasegawa, M., Inohara, N. and Núñez, G. (2006). MEX is a testis-specific E3 ubiquitin ligase that promotes death receptor-induced apoptosis. Biochem. J. 396, 411–417.

Noguchi, T., Lenartowska, M., Rogat, A. D., Frank, D. J. and Miller, K. G. (2008). Proper cellular reorganization during drosophila spermatid individualization depends on actin structures composed of two domains, bundles and meshwork, that are differentially regulated and have different functions. Mol. Biol. Cell 19, 2363–2372.

Pathak, N., Obara, T., Mangos, S., Liu, Y. and Drummond, I. A. (2007). The zebrafish fleer gene encodes an essential regulator of cilia tubulin polyglutamylation. Mol. Biol. Cell 18, 4353–4364.

Politi, Y., Gal, L., Kalifa, Y., Ravid, L., Elazar, Z. and Arama, E. (2014). Paternal mitochondrial destruction after fertilization is mediated by a common endocytic and autophagic pathway in drosophila. Developmental Cell 29, 305–320.

Rajapurohitam, V., Bedard, N. and Wing, S. S. (2002). Control of ubiquitination of proteins in rat tissues by ubiquitin conjugating enzymes and isopeptidases. American Journal of Physiology - Endocrinology and Metabolism 282, E739–E745.

Raunser, S. and Gatsogiannis, C. (2015). Deciphering the tubulin code. Cell 161, 960–961.

Richburg, J. H., Myers, J. L. and Bratton, S. B. (2014). The role of E3 ligases in the ubiquitin-dependent regulation of spermatogenesis. Semin. Cell Dev. Biol. 30, 27–35.

Rogowski, K., Juge, F., van Dijk, J., Wloga, D., Strub, J. M., Levilliers, N., Thomas, D., Bre, M. H., Van Dorsselaer, A., Gaertig, J. et al. (2009a). Evolutionary divergence of enzymatic mechanisms for posttranslational polyglycylation. Cell 137, 1076–1087.

Rogowski, K., Juge, F., van Dijk, J., Wloga, D., Strub, J. M., Levilliers, N., Thomas, D., Bre, M. H., Van Dorsselaer, A., Gaertig, J. et al. (2009b). Evolutionary divergence of enzymatic mechanisms for posttranslational polyglycylation. Cell 137, 1076–1087.

Samji, T., Hong, S. and Means, R. E. (2014). The membrane associated RING-CH proteins: A family of E3 ligases with diverse roles through the cell. International Scholarly Research Notices 2014, 23.

Suryavanshi, S., Edde, B., Fox, L. A., Guerrero, S., Hard, R., Hennessey, T., Kabi, A., Malison, D., Pennock, D., Sale, W. S. et al. (2010). Tubulin glutamylation regulates ciliary motility by altering inner dynein arm activity. Curr. Biol. 20, 435–440.

Tokuyasu, K. T., Peacock, W. J. and Hardy, R. W. (1977). Dynamics of spermiogenesis in drosophila melanogaster. VII. effects of segregation distorter (SD) chromosome. Journal of Ultrastructure Research 58, 96–107.

Tokuyasu, K. T., Peacock, W. J. and Hardy, R. W. (1972). Dynamics of spermiogenesis in drosophila melanogaster. I. individualization process. Z. Zellforsch. Mikrosk. Anat. 124, 479–506.

Venken, K. J., Schulze, K. L., Haelterman, N. A., Pan, H., He, Y., Evans-Holm, M., Carlson, J. W., Levis, R. W., Spradling, A. C., Hoskins, R. A. et al. (2011). MiMIC: A highly versatile transposon insertion resource for engineering drosophila melanogaster genes. Nat. Methods 8, 737–743.

Wang, S., Zheng, H., Esaki, Y., Kelly, F. and Yan, W. (2006). Cullin3 is a KLHL10-interacting protein preferentially expressed during late spermiogenesis. Biol. Reprod. 74, 102–108.

Wei, H., Rollins, J., Fabian, L., Hayes, M., Polevoy, G., Bazinet, C. and Brill, J. A. (2008). Depletion of plasma membrane PtdIns(4,5)P(2) reveals essential roles for phosphoinositides in flagellar biogenesis. Journal of Cell Science 1076–1084.

Wloga, D., Webster, D., Rogowski, K., Bré, M., Levilliers, N., Jerka-Dziadosz, M., Janke, C., Dougan, S. and Gaertig, J. (2009). TTLL3 is a tubulin glycine ligase that regulates the assembly of cilia *dev cell. 2009 jun; 16(6):867-76*. Dev Cell. 16, 867–876.

Wloga, D. and Gaertig, J. (2010). Post-translational modifications of microtubules. J. Cell. Sci. 123, 3447–3455.

Yang, W. L., Zhang, X. and Lin, H. K. (2010). Emerging role of lys-63 ubiquitination in protein kinase and phosphatase activation and cancer development. Oncogene 29, 4493–4503.

Yin, Y., Lin, C., Kim, S. T., Roig, I., Chen, H., Liu, L., Veith, G. M., Jin, R. U., Keeney, S., Jasin, M. et al. (2011). The E3 ubiquitin ligase cullin 4A regulates meiotic progression in mouse spermatogenesis. Dev. Biol. 356, 51–62.

Yogo, K., Tojima, H., Ohno, J. Y., Ogawa, T., Nakamura, N., Hirose, S., Takeya, T. and Kohsaka, T. (2012). Identification of SAMT family proteins as substrates of MARCH11 in mouse spermatids. Histochem. Cell Biol. 137, 53–65.

Zhang, P. and Spradling, A. C. (1993). Efficient and dispersed local P element transposition from drosophila females. Genetics 133, 361–373.

Zhao, B., Ito, K., Iyengar, P. V., Hirose, S. and Nakamura, N. (2013). MARCH7 E3 ubiquitin ligase is highly expressed in developing spermatids of rats and its possible involvement in head and tail formation. Histochem. Cell Biol. 139, 447–460.

Zhi, X., Zhao, D., Wang, Z., Zhou, Z., Wang, C., Chen, W., Liu, R. and Chen, C. (2013). E3 ubiquitin ligase RNF126 promotes cancer cell proliferation by targeting the tumor suppressor p21 for ubiquitin-mediated degradation. Cancer Res. 73, 385–394.

